# 3D genome architecture regulates the traffic of transcription factors throughout human chromosomes

**DOI:** 10.64898/2026.01.06.698019

**Authors:** Amit Das, Ryan R. Cheng, Davit A. Potoyan, Michele Di Pierro

## Abstract

Transcription factors (TFs) control the rate of transcription of genetic information by binding to specific DNA sequences. The time needed for a TF to find its specific target sites is a bottleneck to the genetic response mechanism. While TF target site search is a well-studied problem, the effect of genome 3D architecture on the TF target search times is poorly understood. Here, we use accurate and cell-specific 3D structural ensembles of human chromosomes to investigate how the spatial organization of binding sites on chromosomes influences the dynamics of TFs. We use Chromatin Immuno-Precipitation data to map the position of binding sites for several TFs on chromosomal structures and simulate the dynamics of individual TF within chromosomal territories. We find that the distribution of binding sites along chromosomes cooperates with the 3D folding of the chromatin fiber to induce dynamics in which TFs tend to visit sites distributed sequentially along the genome. In this way, genome 3D architecture appears to reduce the time each TF spends in the unbound state while commuting from one target site to the other. At the same time, genome 3D architecture further reduces the flux of TFs between binding sites already well separated along the genome, effectively isolating distant clusters of binding sites. We compare the TF traffic patterns generated by the 3D structures of human chromosomes with those generated by several alternative structural models characterized by increasing randomness. Finally, we study the effect of lengthwise compaction and phase separation, known architectural features of the human genome, in TFs target search. In short, our analysis demonstrates that genome architecture regulates the traffic of TFs within chromosomal territories and reduces the time each TF spends commuting between binding sites.

## A. Significance

Transcription factors must efficiently locate multiple regulatory targets within the crowded nucleus. By combining realistic 3D chromosome structures with genome-wide binding maps, we show that genome folding organizes transcription factor motion into modular, neighborhood-level exploration, enhancing intra-neighborhood diffusion while insulating distant regulatory programs. These findings uncover a previously unrecognized dynamical role of genome architecture beyond spatial proximity.

## B. Introduction

Transcription is the first step among the series of processes involved in the expression of a gene. Transcription factors (TFs) are proteins that bind to specific DNA sequences and regulate gene expression by controlling the rate of transcription initiation for one or more genes. In eukaryotes, TFs have evolved a rich repertoire of molecular mechanisms to control the process of transcription [1]. To accomplish its function, a TF must first find one of its specific target sites along the genome. Because of this, the time associated with TFs target site search plays a fundamental role in the genetic response mechanism; search time is likely subject to biological regulation and evolutionary dynamics.

A single TF typically binds to a large number of DNA regulatory sequences [2] which are found and bound by a similarly large number of TF copies diffusing throughout the nucleus. Facilitated diffusion, a combination of 1D sliding along the genome and diffusion in 3D, has been proposed as a mechanism to achieve more efficient passive transport of DNAbinding proteins compared to 3D diffusion alone. [3–8].

Polymer models for chromatin have been used to study the effect of chromatin density and epigenetic marks over the diffusion of DNA-binding proteins [9–11]. Standard and facilitated diffusion models have been coupled with simplified models of chromatin to study the effect of the 3D organization on the process of target site search. [9–15]. These early attempts at studying the repercussion of nuclear architecture on the target search problem were stymied, however, by the lack of many of the essential features of 3D genome organization and quantitative data on the 1D positioning of the binding sites along the genome. Due to the explosion of activity to characterize genome architecture seen in recent years, we can now introduce accurate 1D positioning of binding sites and 3D folding of the fiber into a single theoretical model for TF dynamics in the nuclear environment.

We investigate how the traffic of TFs diffusing throughout human chromosomes is influenced by the spatial organization of binding sites on chromosomes. Because of the scarcity of information about co-factors, kinetics, and molecular mechanisms of TFs binding and 1D sliding (if any), we focus exclusively on the effects architecture exerts over the 3D diffusion of the molecules in solution. Similar to how urban architecture evolves to reduce citizens’ commute times, we find that the distribution of binding sites along chromosomes cooperates with the 3D folding of the chromatin fiber to reduce the time TFs spend in solution, commuting from one binding site to another. While genome architecture influences the motion of DNA-binding proteins moving throughout nuclei, proteins also shape the DNA structure through self-assembly and phase separation. [16–21]. The feedback loop between genome architecture and DNA-binding proteins is, however, characterized by a strong separation of timescales, with the 3D distribution of small, fast-moving TFs rearranging much faster than chromosome. We exploit this separation of timescales to decouple the two problems; in our simulations, we use detailed yet static chromosomal configurations to study TFs pathways of diffusion within them.

The structural ensembles of chromosomes are fluid and yet specific to cell type and phase of life [22–24]. In the last few years, our understanding of these ensembles has improved substantially, mainly as a result of the development of new experimental techniques, such as DNA-DNA proximity ligation assays [25] and super-resolution microscopy [26, 27]. DNA proximity ligation assays have shown that the organization of interphase nuclei and chromosomes in mammals is dominated by the segregation of chromatin into genomic compartments [25, 28, 29]. Additionally, smaller contact domains exist within compartments, known as topologically associating domains (TADs) [25, 28–30]. DNA sequences recruiting the architectural protein CTCF often mark the boundaries of TADs [29, 31]; at these loci, CTCF binding motifs are largely found in a convergent orientation, i.e. pointing toward each other with the intervening DNA forming a loop. To explain this convergent orientation, it has been proposed that these loops form through a process of chromatin extrusion [32–35]. In this model, structural maintenance of chromosome (SMC) complexes, which are ring-shaped protein assemblies, extrude a DNA loop until they are stopped by CTCF molecules bound to an inward-pointing motif. The Minimal Chromatin Model (MiChroM) is a physical framework that, in light of the experimental observations just outlined, explains chromosome organization through the processes of lengthwise compaction and liquid phase separation [36, 37]. According to MiChroM, lengthwise compaction results from the activities of DNA-extruding SMC complexes [34, 38] and possibly other motors, while epigenetically-regulated liquid phase separation is instead responsible for chromatin compartmentalization.

The genomic conformations of human chromosomes calculated using the MiChroM are cell type-specific and in remarkable agreement with data from both DNA-DNA proximity ligation assays and 3D fluorescence microscopy [22, 24, 37, 39]. In this work, these data-driven 3D structural ensembles are the starting point to study the dynamics of TFs in a realistic and cell-specific nuclear environment. We focus on a few different TFs with important roles in the regulation of core biological processes and diseases. We study how TFs diffuse within the chromosomal territory in the presence of their own specific 3D spatial distribution of binding sites along DNA. To dissect the effects of motor activity and epigenetics on TF traffic regulation, we finally consider multiple descriptions of chromatin organization; starting from the full architecture and gradually removing the various organizing features such as phase separation, lengthwise compaction, and the 1D positioning of target sites along the genome. These analyses bring out regulatory effects associated to each of the layers of 3D genome organization.

## C. A physical framework to study the dynamics of TFs in realistic chromosomal environments

To understand the effects of the 3D genome architecture on the dynamics of TFs we need accurate and cell-type-specific 3D conformations of chromosomes and the binding energy landscape of the TFs. Ensembles of chromosomal conformations have been calculated using the MiChroM theory for the autosomes of a variety of human cell lines [24] [36] [37] and are accessible through the Nucleome Data Bank (NDB) [39]. The MiChroM chromosomal structural ensembles have been shown to predict accurately the results of experiments of both DNA-DNA proximity ligation and 3D fluorescence microscopy. The binding energy landscape of TFs can instead be extracted form ChIP-seq data available from the ENCODE database [40, 41] (See Methods for more details)

For each chromosome, we map the binding energies Δ*G*_*b*_ on the conformations of chromatin obtained from simulation, thus obtaining the 3D distribution of the binding sites (see *Methods*). We then carry on molecular dynamics simulations (details in *Methods*) in which TFs can diffuse throughout chromosomal territories as well as bind and unbind target sites according to their respective free energies. Individual TFs are represented by spherical particles of appropriate size that, when unbound, move according to Langevin dynamics. The binding and unbinding of a TF molecule to a target site *i* is modeled with a potential well of depth Δ*G*_*i*_ which extends from the position *r*_*i*_ of the binding site outward over the sphere delimited by a radius *r*_*b*_. Because our interest lies in studying the way TFs diffuse through chromosomes (i.e., in the unbound state exclusively) the details of the binding and unbinding chemical kinetics are not considered in this study. Besides binding to target sites, no additional interactions are considered between chromatin and TFs. In particular, no steric interactions are considered; while this is undoubtedly a simplification, this is consistent with the established notion that TFs can effectively penetrate chromatin to reach anywhere in the nucleus [42–44]. By comparing chromatin structural models with similar, approximately uniform levels of crowding, we effectively control for the crowding-induced slowdown of three-dimensional diffusion. More details on the simulation protocols are provided in the Methods. We study the dynamics of several TFs. A majority of our results involve the REL-associated protein (RELA) from the well-studied Nuclear Factor Kappa B (NF-κB) family of transcription factors. RELA, also known as p65, has been identified as an important factor in the development of the immune system and in oncogenesis [2] [42] [45]. We compare the results for RELA with those obtained for MAZ (MYC-associated Zinc finger protein), a TF involved in the activation MYC family of oncogenes related to carcinomas in the cervix, breast, colon, lung, and stomach. Additionally, we study the dynamics of two pioneer TFs [46]: POU5F1 (also called OCT-4) and FOXA1. POU5F1 is a involved in stem cell pluripotency [47], while FOXA1 is active during cell differentiation of the liver and pancreas [48, 49] as well as implicated in breast cancer [50].

## D. Genome architecture facilitates fast TF commute times

We analyze individual TF trajectories to identify the binding and unbinding events to and from the chromatin fiber and convert each trajectory into a sequence of intervals of time spent in bound and unbound states (Fig. 2b). The bound intervals of time are the residence times (τ_*res*_) of the TFs on chromosomes, while the intervals between successive binding events are the waiting times τ_*w*_, or commute times. The distribution of the residence times τ_*res*_ is roughly bi-exponential (Figure S1) and independent from chromosomal architecture. This bi-exponential distribution reflects the hundreds of individual binding energies extracted from ChIP-seq and the simple kinetic model we assumed for binding and unbinding. Despite the crudeness of our assumptions regarding binding kinetics, our simulated residence times are nevertheless in good agreement with experiments measuring the DNA residence times of TFs in vivo [4, 51, 52].

**FIG. 1.**
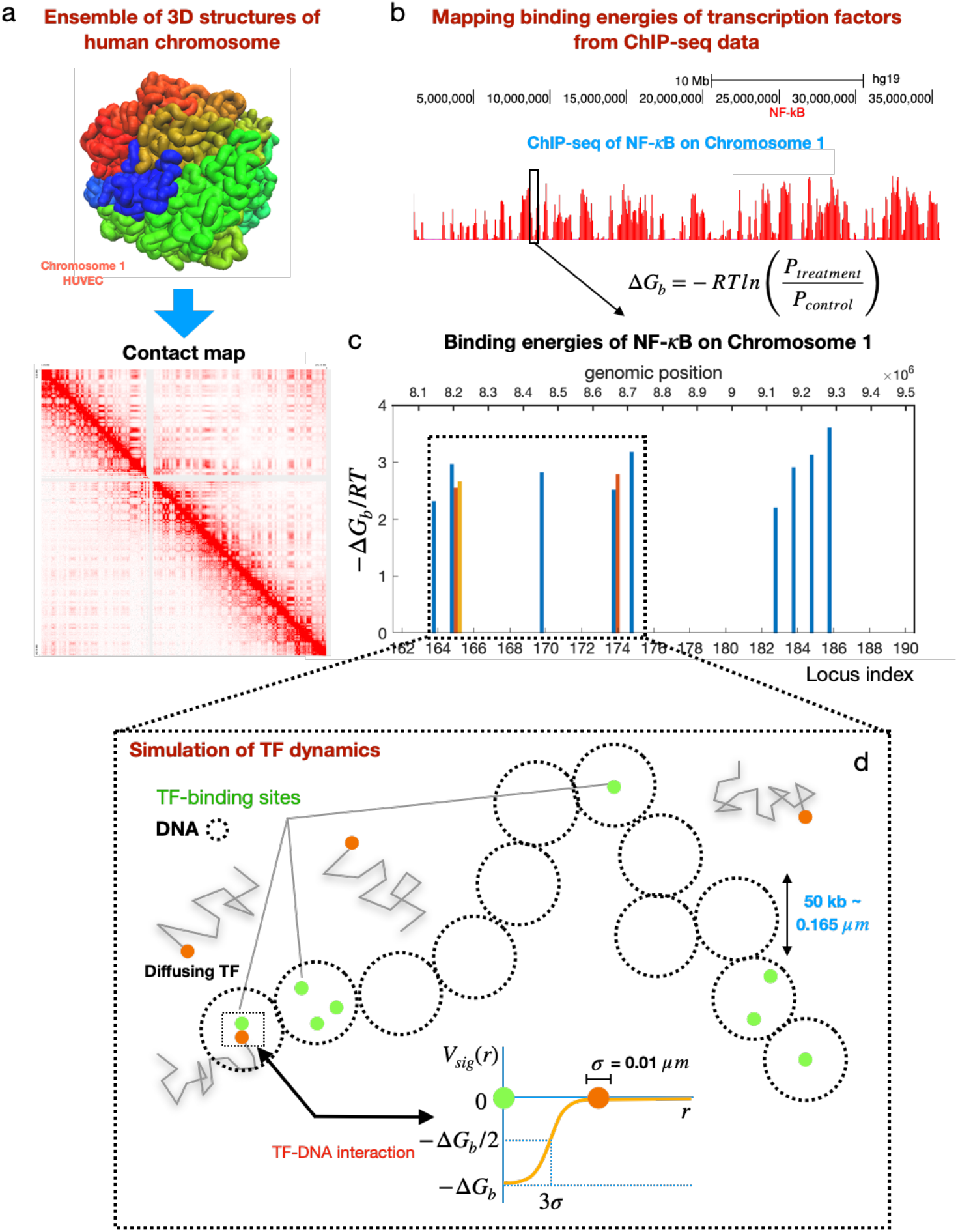
Workflow for simulation of transcription factor (TF) dynamics using ensemble of human chromosome structures. (a) Ensemble of accurate 3D structures of human chromosomes, one representative chromosome structure shown in top, which fully recapitulates the Hi-C chromatin contact map, example shown in bottom). (b) Mapping of binding energies for TF-DNA binding sites from ChIP-seq data for specific TF’s available through the ENCODE database. (c) Mapped binding energies are shown for the transcription factor NF-κB for a small set of binding sites on chromosome 1. (d) Schematic of our simulation of TF dynamics on human chromosomes. A segment of chromosome is shown with overlay of binding sites. In our simulations we just have the binding sites that interact with the TFs with a sigmoid interaction potential *V*_*sig*_(*r*) where *r* is the distance between a diffusing TF and a DNA binding site and strength Δ*G*_*b*_, the free energy of binding.

**FIG. 2.**
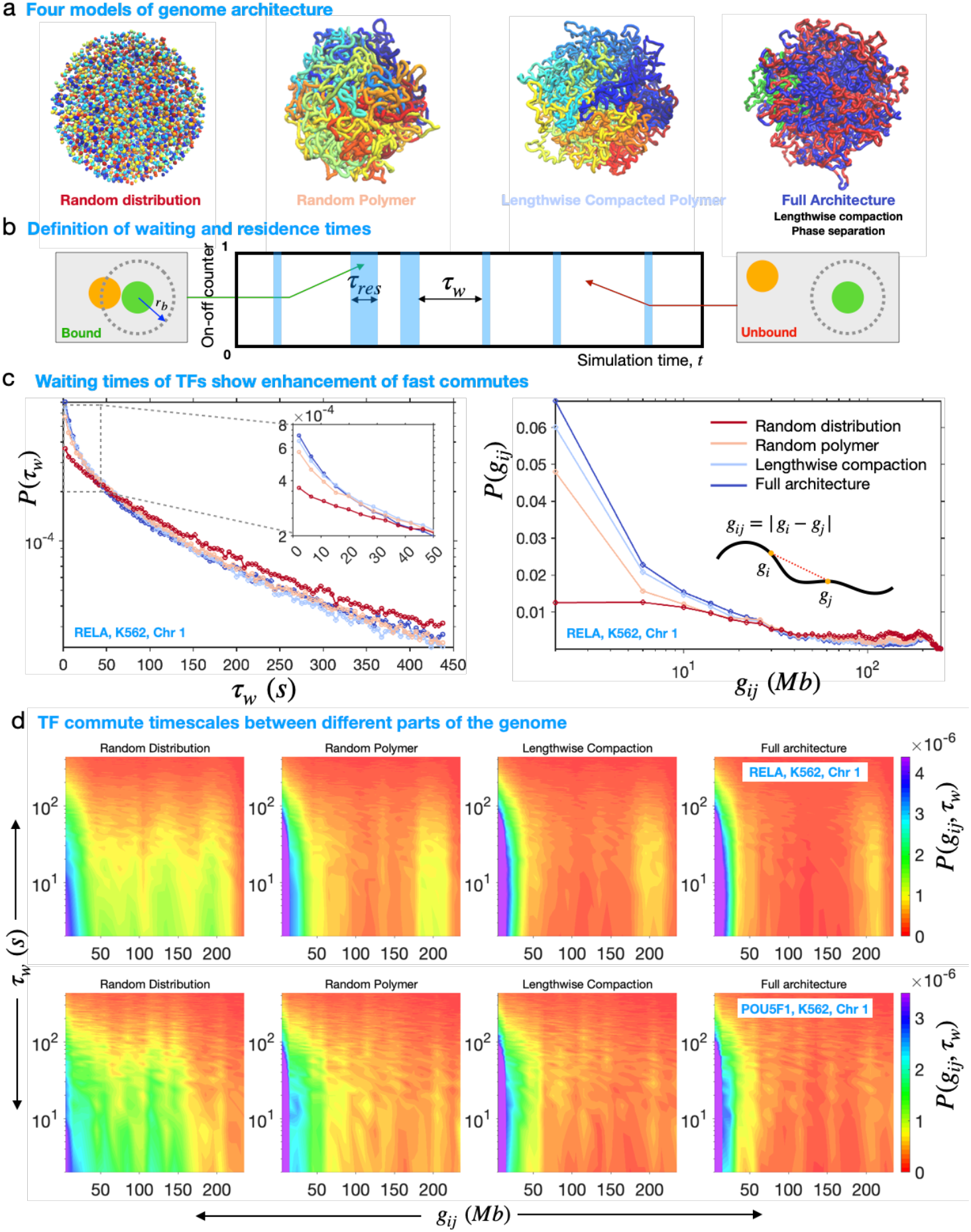
Four chromatin models reveal hierarchical speed-up of TF on-off dynamics on DNA by different features of chromatin organization. (a) Comparison of the 3D structure of four chromatin models, illustrated for human chromosome 1. Tube representation are shown for the polymer models, while Van der Waal’s space filling representation is shown for the random organization. The coloring method for the full 3D architecture (first from right) is highlighting the chemical identities of chromatin beads, while that for others is the end to end index of chromatin beads. Illustrations made using VMD [68]. (b) Schematic showing the method of determining the on-off state of TFs (top). Here *r*_*b*_ is a cutoff radius. Any TF molecule (chrome) found within *r*_*b*_ from a TF binding site (green) on chromatin is assigned to be bound and considered unbound otherwise. A representative time series of on-off dynamics of a TF is shown in the bottom panel. (c) Full probability distribution functions (PDFs) of commute times (*t*_*w*_) (left panel) and genomic distances (*g*_*i j*_ ) corresponding to the commuting events of RELA molecules observed for the different models of organization in chromosome 1 from K562 cells. Any given PDF is generated pooling together the waiting times or genomic distances extracted from all the individual trajectories of TF molecules collected from independent simulations on at least 10000 or more chromosome structures. (d) Bi-variate distributions of τ_*w*_ and *g*_*i j*_, given by *P*(*g*_*i j*_, τ_*w*_) as heat-maps for RELA (top row) and POU5F1 (bottom row). For both the TFs, *P*(*g*_*i j*_, τ_*w*_) are shown for all the four models of chromosome 1 from K562 cells.

The distribution of waiting times, however, exhibits a clear dependence on chromosomal conformation. To study the effect of genome architecture on the probability density function (PDF) of commute times, we compare the results obtained with the data-driven, detailed, structural ensembles of human chromosomes with several simpler models of chromosomes characterized by increasing levels of the organization.

The crudest model is a spherical uniform random distribution of the target binding sites. Here, there is no structural organization and the only relevant parameter is the density of binding sites, which is set to reproduce the one observed in the data-driven ensembles [23, 36] and imaging experiments [53– 55]. This constant average density of target binding sites characterizes all the chromosome models studied.

The second model is a self-avoiding random polymer that preserves the positioning of the target binding sites along the chromatin fiber. This model already includes a significant amount of organization, as the positioning of the DNA binding sites of a specific TF, which we extract from ChIP-seq assays, is far from random. Binding sites are typically more frequently found in active chromatin (the A compartment) and there distributed in clusters inter-spaced by longer stretches of DNA devoid of any binding sites.

Additional organizational information beyond the 1D organization of genetic elements (above) is contained within the 3D folding of chromatin, which determines the spatial organization of the target binding sites. The MiChroM Hamiltonian, which we use to recapitulate the 3D folding of human chromosomes, relies on two energy terms: phase separation and lengthwise compaction. Aggregation of chromatin characterized by similar epigenetic marking patterns leads to the formation of microphases within chromosomes, which segregate genes in space and result in the compartmental interaction patterns observed in proximity ligation assays. Phase separation correlates strongly with particular combinations of histone modifications [37]; as such, genomic interactions due to phase separation depend on the chromosome identity and cell type. On the contrary, lengthwise compaction appears to be largely similar among chromosomes and cell types within an organism, albeit changing dramatically along the cell cycle and among different organisms. The origin of lengthwise compaction appears to lie in the activity of DNA-extruding complexes, such as those belonging to the SMC family and other motors [38, 56]. A lengthwise compacted polymer is the third structural model we compare with the complete structural ensembles of chromosomes, thus separating the contribution of motor activity on regulating the traffic of TFs from that of phase separation.

In increasing order of organization, the four models of chromosomes considered are then I) a random distribution of the experimentally determined DNA binding sites, II) the same binding sites distributed according to their genomic positioning along a random self-avoiding polymer, III) the binding sites distributed according to their genomic positioning along a lengthwise compacted polymer, and IV) the binding sites distributed according to their genomic positioning along a lengthwise compacted and phase separated polymer, which reproduces the chromosomal structural ensemble observed experimentally in human cells.

Each of our simulations are ∼2200 seconds long when converted into real units. We rarely see any commute times longer than ~450 seconds, and hence, restrict our analysis to timescales less than ∼450 seconds. For each of the four models, the PDF of commute times, *P*(τ_*w*_) is multi-exponential with a fast component and 2-3 slow components (Fig. 2c-left). Strikingly, genome architecture appears to favor local interactions, enhancing the probability of short commutes. This effect increases monotonically with each layer of organizational complexity and reaches its maximum for the data-driven chromosomal ensembles (Fig. 2c-left). This reflects a separation of time-scales associated with distinct modes of motion of TFs across chromosomes. The fast component reflects jumps between binding sites that are spatially close, including rebinding events. In our city analogy, these short commutes could be described as trips within the same neighborhood. On the other hand, the slow components arise from prolonged periods in which TFs remain unbound while diffusing in between distant binding sites. In our analogy this corresponds to commutes between different neighborhoods.

Clearly, the amount of time the TFs spend in solution versus the time bound to their target sites depends on the 3D organization of the binding sites, with each layer of structural organization contributing to reducing the TF commute times. The least efficient distribution of binding sites−the one with the lowest probability of shortest commute times−is found to be the uniform distribution. Next follows the 3D distribution associated with the random polymer. The speed-up of TF dynamics due to the 1D positioning of binding sites along DNA is the most significant, asserting the longstanding notion that the positioning of genes has consequential effects and it is far from random. Both lengthwise compaction and phase separation further reduce the average commute times between targets. Surprisingly, the speed-up associated with lengthwise compaction is larger than the one associated with phase separation, suggesting a new functional effect for DNA-extrusion. The same results were found consistently across chromosomes and for other TFs (Fig. S2).

To confirm the previous assertion that 1D positioning of binding sites along the DNA chain is responsible for the speed-up, we look at the PDFs of the commuting genomic distances associated with all the TF commutes. These PDFs corroborate the PDFs of the commute times. We see a similar enhancement of the probability of short genomic distances from random distribution to genome architecture (Fig. 2cright). This constitutes another piece of evidence supporting the 3D organization of the genome in enhancing within-neighborhood TF commutes.

Next, we check how the TF commutes between different chromatin neighborhoods originate from fast and slow commutes, respectively. To understand this, we consider a bivariate PDF of the commute times and the genomic distances. Figure 2d shows these PDFs as heatmaps for two different TFs on chromosome 1 from the K562 cell type. First, we consider the results for RELA (top row). Among the four models, the random distribution reflects the weakest correspondence between commute times and genomic distances. The polymer models, however, are drastically different. The random polymer shows a depletion of the bivariate distribution at genomic distances in the range ≈50− 160 Mb for fast to intermediate commute times (up to 200 s). This depletion is more pronounced for the lengthwise compacted polymer and strongest for the full genome architecture. Similar trends hold true for another TF called POU5F1, also known as Oct4, as shown by PDFs in Fig. 2d-bottom row.

This analysis reveals a strong regulatory influence of the different organizational features of the genome on the TF commute times. The gradual depletion of the bi-variate PDF at intermediate genomic distances from random polymer to full genome architecture means that fast TF commutes can occur on the genome if the genomic distances are reasonably small (< 50 Mb) or even quite large as hundreds of Megabasepairs (> 166 Mb). In the next section we will look at the PDFs of Cartesian distances corresponding to genomic distances defined by these specific ranges to understand this depletion effect and how it depends on the spatial positioning of the TF binding sites in 3D.

## E. Interplay of 3D genome architecture and sequence organizes TF traffic into intra- and inter-compartment commutes

We measure the Cartesian jump lengths *r*_*ij*_ on the chromosome corresponding to the commuting events of the TFs, expressed as the Euclidean distance between subsequently visited binding sites *i* and *j*. To identify the jump lengths corresponding to different chromatin neighborhoods, we focus on three different ranges of genomic distance, *g*_*ij*_: small: *g*_*ij*_ ≤ 50 *Mb*; intermediate: 50 *Mb < g*_*ij*_ ≤ 166 *Mb* and large:*g*_*ij*_ *>* 166 *Mb*. Here, the choice of 166 Mb is specifically made based on the evidence in Fig. 2d for the full architecture model of Chromosome 1. Incidentally, 166 *Mb* is also equivalent to 2/3-rd of the total chromosome length in base pairs. The PDFs of *r*_*ij*_ for RELA (Fig. 3a-top row) in the small genomic range show a substantial enhancement of short Cartesian jumps by the genome architecture. A strong peak appears near *r*_*ij*_ ≈ 200 nm, which is close to the spatial resolution (bead diameter) in MiChroM [24]. Here, the peak captures the short-range hops and indicates a strong enrichment of these events.

**FIG. 3.**
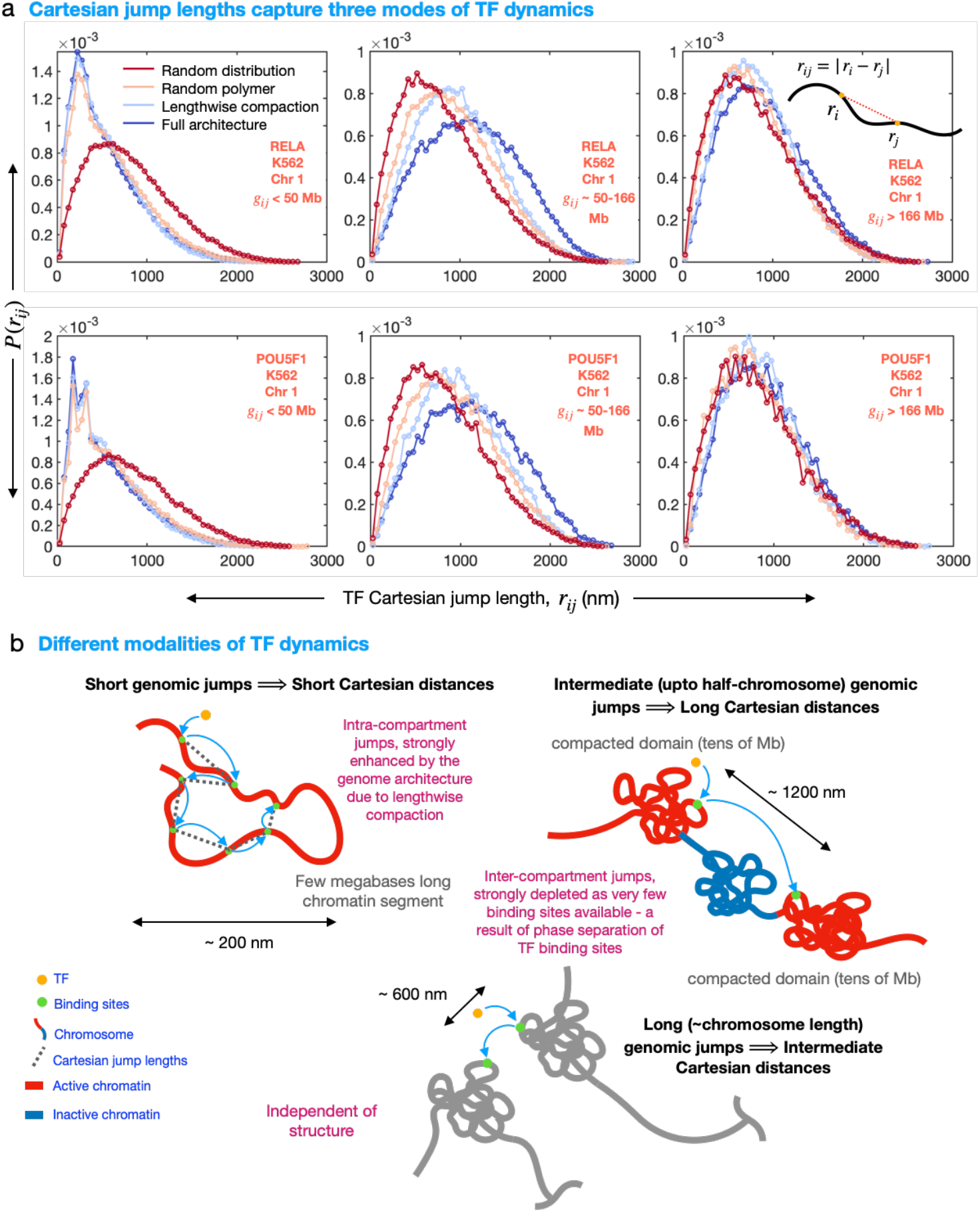
Spatial commute distances reveal intra- and inter-compartmental commutes of TFs. (a) PDFs of Cartesian jump distances for three different zones of genomic distances: *g*_*i j*_ ≤ 50 *Mb* (left), 50 *< g*_*i j*_ ≤ 166 *Mb* (middle) and *g*_*i j*_ ≥ 166 *Mb* (right). Here, 166 *Mb* is ≈2*/*3-rd of chromosome 1. Results are shown for RELA molecules (top row) and POU5F1 molecules (bottom row) exploring target sites on chromosome 1 from K562 cells. (b) Schematic representation of the various modes of TF dynamics on chromatin, illustrating the types of genomic jump distances that could lead to three specific types of Cartesian jump lengths that are most probable as revealed by plots in (a). These cartoons provide illustrations for intra-compartmental commutes (top left), inter-compartmental commutes that are far away and separated by inactive chromatin (top right) and inter-compartmental commutes that are proximal owing to lenghtwise compaction (bottom). The first and final kind of commutes are promoted by the genome architecture whereas the commutes of the second kind are suppressed.

The intermediate genomic range reveals a different behavior. Here, the order of the peaks switches entirely, and also the strength of the peak diminishes gradually from the random distribution model to the entire architecture model. This is reminiscent of the depletion of TF commutes in this range seen before in Fig. 2d. *P*(*r*_*ij*_) from the four models appear indistinguishable for the large genomic range. We find similar evidence for POU5F1 (Fig. 3a-bottom row) and for different chromosomes (Fig. S4).

These results are recapitulated schematically in Fig. 3b. In short, genome architecture generates a strong enrichment of short commutes at short genomic range, which correspond to spatial distances ∼ 200 *nm*. These jumps arise from the target sites separated by hundreds of kilobases to a few megabases, which possibly lie in the same chromatin compartment. At the same time, genome architecture determines a substantial depletion of short commutes in the intermediate genomic range, which encompasses up to ∼ 2*/*3 of the chromosome length and measures up to more than a micron. At large genomic distance, TFs commutes appear to be insensitive with respect to spatial organization.

Our findings reveal an interplay between 3D genome architecture and the linear arrangement of transcription factor binding sites along the chromatin fiber. This interaction regulates TFs movements, facilitating intra-neighborhood traffic while restricting inter-neighborhood communications.

## F. Discussion and conclusion

The exploration of genome organization in 3D is a major achievement of the last decade, which paves the way to understanding how the different binding sites targeted by diffusing DNA-binding proteins are distributed within the nucleus. This information is a key component in determining the dynamics of transcription factors across chromosomes; a component that, previously, could not be analyzed. We used an ensemble of accurate, data-driven, spatial distributions of target sites to explore their effects on the search process. We focused on a few transcription factors characterized by a large number of DNA binding sites, and repeated the analysis for all chromosomes in several immortalized human cell lines. The first mechanism influencing the distribution of target sites within nuclei is their positioning along the chromatin polymer. Genome architecture, intended as the folding in 3D of such a polymer, contributes two additional organizing mechanisms: lengthwise compaction of the polymer and phase separation. We show that all three organizing mechanisms matter. The interplay between 1D positioning of binding site along chromosomes and 3D folding of the chromosomes regulates the traffic of transcription factors as they hop from one binding site to another. Short commutes between neighboring binding sites within individual clusters are enhanced by the underlying genome architecture. Instead, commutes in the intermediate range, among binding sites belonging to neighboring clusters, appear to be suppressed. Finally, infrequent, long commutes between distant clusters of binding sites appear unaffected by genome architecture. Overall, the regulatory effect on TFs dynamics seems to favor a modularization of the genetic response, enhancing communication within modules while insulating them from one another. While our results clearly delineate how genome architecture gives rise to these effects, our analysis does not yet establish their functional relevance. With accelerating progress in characterizing genome 3D architecture, we expect the next few years to dramatically advance our understanding of the physical genome, intended as the complex set of large macromolecules moving and interacting in three dimensions within cell nuclei. Conversely, a clear understanding of the physical processes underlying the motion and operation of genomes will greatly facilitate the resolution of long-standing questions in biology, such as gene regulation.

## METHODS

### G. Derivation of TF-DNA binding free energy from ChIP-seq data

In this section, we derive the free energy of interaction between a transcription factor (TF) with DNA, and describe how we can estimate it from the ChIP-seq data. ChIP-seq experiments typically consists in two assays performed on chromatin: one in which the DNA sequences bound by a specific target protein are enriched for through immuno-precipitation (signal) and one where no selection of DNA sequences is made (control). Measurements typically report the relative enhancements of the population of the DNA sequences bound by the target protein with respect to the control population.

In spirit of the formulations in Refs [57–59], we start by considering the following reversible reaction between a TF and a short segment of DNA, represented by sequence 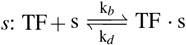 where *k*_*b*_ and *k*_*d*_ are the rates of binding and dissociation of the TF, respectively, which depend on the DNA sequence *s*. At equilibrium, the probability of finding the TF bound to sequence *s* is given by

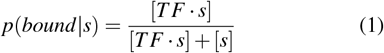

where square brackets represent the molar concentrations of the respective species. Replacing with the equilibrium binding constant *K*(*s*) = *k*_*b*_*/k*_*d*_ = [*TF · s*]*/*([*TF*][*s*]), we get

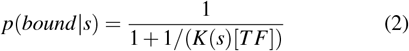

Using the definition of chemical potential of TF: *µ*_*TF*_ = *RTln*[*TF*] and equilibrium binding free energy: Δ*G*(*s*) = −*RTlnK*(*s*), Eq. 2 becomes

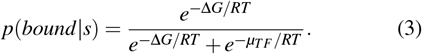

In a typical ChIP-seq assay there are many such DNA sequences that interact with the TF. In a given set of sequences *s*_*i*_ that compete for binding to the TF, the probability that any sequence *s*_*i*_ to be found in the bound state can be written using the Bayes’ theorem:

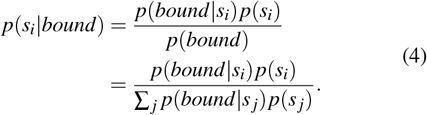

where *p*(*s*_*i*_) represents the probability of any given sequence in the input distribution of sequences. This is obtained from the control experiment of the ChIP-seq assay where the DNA sequences are not stimulated for the binding of the specific target TF or stimulated for a non-specific protein that binds everywhere. We will now use the results of Eq. 3 in a limit [*TF*] *<<* [*DNA*], which renders *µ*_*TF*_→ − ∞ and the denominator a constant. Therefore, from Eq. 4 we find that

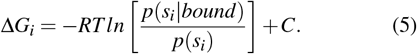

where 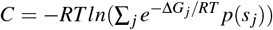 which is essentially a constant. Setting this constant to zero we define the relative binding free energy of the TF interacting with sequence *s*_*i*_, simply referred to as Δ*G*_*b*_ for conciseness from this point onward:

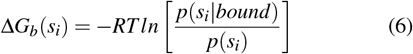

The ratio in the right hand side is obtained from the ChIP-seq peaks. Any peak represents the enrichment of a given stretch of DNA in the population of the TF-bound sequences over the control population. Therefore, we calculate the above ratio by integrating the ChIP-seq peak corresponding to sequence *s*_*i*_. The ChIP-seq tracks are analyzed on Galaxy webserver [40]. Alignments for the control sample and the treatment sample are obtained from the .bam files deposited on the ENCODE database (http://encodeproject.org). Then MACS peak caller [60] version 2 is used to determine the peak positions and their widths. The peaks are integrated using the bedtools program [61]. The used .bam files for the studied TFs are mentioned below: For RELA, we have used ENCFF954LMM (control) and ENCFF392CDE (treatment). For POUF51, we have used ENCFF841BOW (control) and ENCFF190HSK (treatment). Both data sets are aligned with the hg19 assembly of human genome.

### H. Simulation details

We use a 5 *µ*m cubic box with periodic boundaries in all of *x, y* and *z* directions for our simulations of TF dynamics around a human chromosome structure made of spherical particles representing TF binding sites. A representative chromatin segment simulated is shown schematically in Fig. 1d. We perform every simulation with the same concentration of TFs (500 copies per simulation box), which catpures the observed range of TF concentrations in mammalian cell (typically 10^3^ − 10^6^ copies per cell) [62]. Every simulation contains one single chromosome placed within the simulation box and the chromosome is surrounded by a uniformly distribution of the TFs. Furthermore, we make an assumption that the chromosome structure evolves much slowly in time compared to the movement of the TFs. So, the chromosomal particles are kept frozen in space throughout any simulation.

TFs are modeled as diffusing spherical particles with a diameter σ ≈ 10 nm. This is the approximate physical size of a typical TF in mammalian cells [63]. For simplicity, we use the same physical size uniformly for all the particles posing as chromosomal binding sites, which describes the hardcore of the TF-DNA interaction represented in our model by the Weeks-Chandler-Anderson (WCA) potential [64]. We represent the attractive part of the TF-DNA interaction by a potential function of sigmoid shape where the depth of the potential is given by the corresponding Δ*G*_*b*_ (shown in Fig. 1d). TFs are weakly interacting via the WCA potential among themselves.

While the chromosomal structural ensembles are characterized by a 50 kb resolution, typical of Hi-C experiments, the binding energy landscape along genome is characterized by a higher resolution typical of ChIP-seq experiments. Because of this, often more than one binding site, i.e. multiple ChIPseq peaks, fall within one 50 kb regions (Fig. 1c). To bridge the gap between the resolution of the two data sets used, for any 50 kb region where *n* such TF binding sites are found, we distribute these *n* sites randomly within the volume occupied by the chromatin segment in the nucleus. Below the resolution accessible through the structural model, a uniform random distribution of binding sites reflects the absence of any additional insight into their spatial arrangement (Fig. 1d). Our simulations involving various models of the chromatin are carried out with a chromatin density comparable to the density reported earlier by the MiChroM model [23, 36] and the imaging experiments [53–55].

TFs diffuse in a medium that represents the viscous background of cell nucleus. We capture this through a Langevin dynamics simulation of the TFs. The effects from the medium are modeled through a frictional drag given by Γ = 6πη_*nu*_σ, where η_*nu*_ is the viscosity of the nucleoplasm. Measured viscosity of the nucleoplasm varies widely [65] based on the technique used. Here we used an intermediate value of η_*nu*_ = 10 Pa-s which is close to that measured using microrheology experiments [66]. We also fix the temperature of the simulation at physiological 310 K, which defines *k*_*B*_*T*, the Boltzmann constant *k*_*B*_ times the absolute temperature, *T* . We use the open-source package HOOMD-Blue [67] to simulate Langevin dynamics of the TFs.

We use a timestep Δ*t* = 0.01τ to integrate the equations of motion of the TFs, where τ = Γσ^2^*/k*_*B*_*T* = 44 ms that defines the unit of time of our simulations. All lengths in our simulation are measured in units of σ and all energy in units of *k*_*B*_*T*.

### I. Analysis of TF trajectories

We analyze the single TF trajectories to identify the binding events and locate the moments it detaches from a site. To do this, we define a binding radius *r*_*b*_ around every binding site. A TF molecule is considered bound if its position falls within a distance *r*_*b*_ from one of its target binding sites and otherwise unbound (Fig. 2b). We apply this condition to all the TFs and extract the sequences of intervals of bound and unbound states from each TF. These sequences are then used to determine the distributions of waiting times and DNA residence times of the TFs.

## J. Acknowledgments

We thank the members of the Nuclear Physics working group in the Center for Theoretical Biological Physics for valuable discussion. This research was supported by the National Science Foundation through the awards PHY-2412651 and PHY-2019745, and by the National Institute of General Medical Sciences of the NIH under award R35-GM146852 (to MDP) and award R35-GM138243 (to DAP). The content is solely the responsibility of the authors and does not necessarily represent the official views of the funding agencies.

## SUPPLEMENTARY DATA

**FIG. S1.**
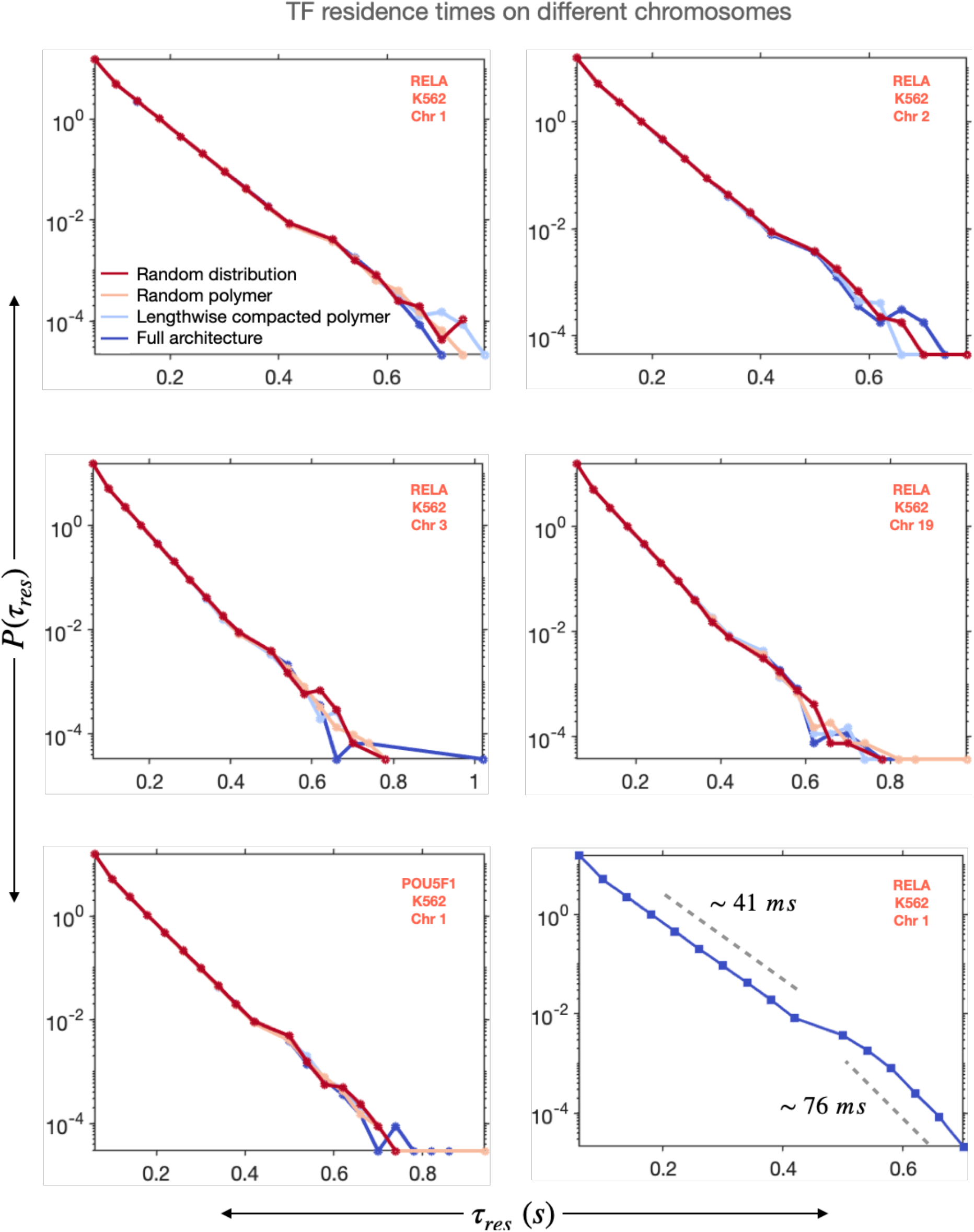
PDFs of TF residence time on different chromosomes.

**FIG. S2.**
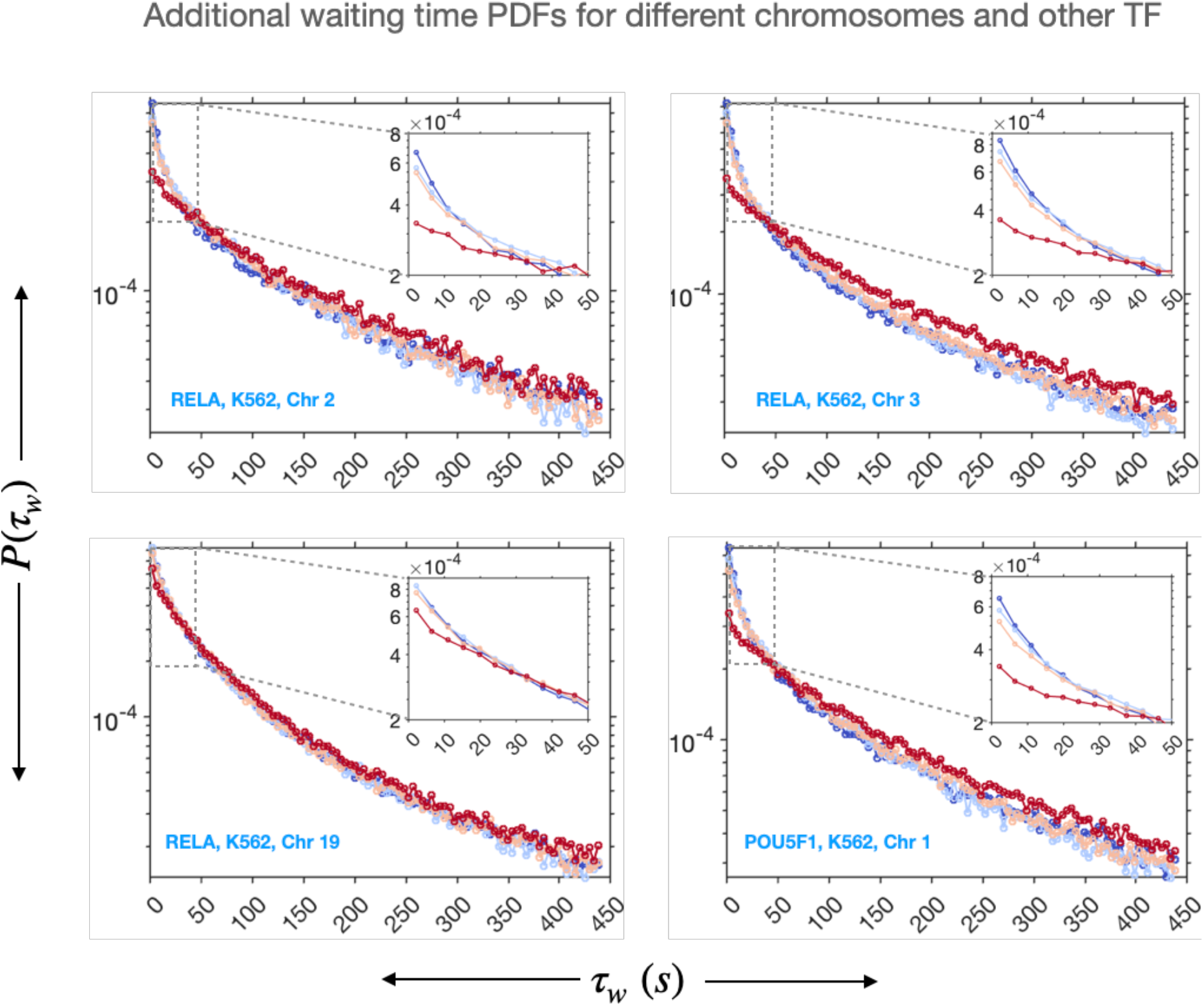
Additional data on different chromosomes for different TFs.

**FIG. S3.**
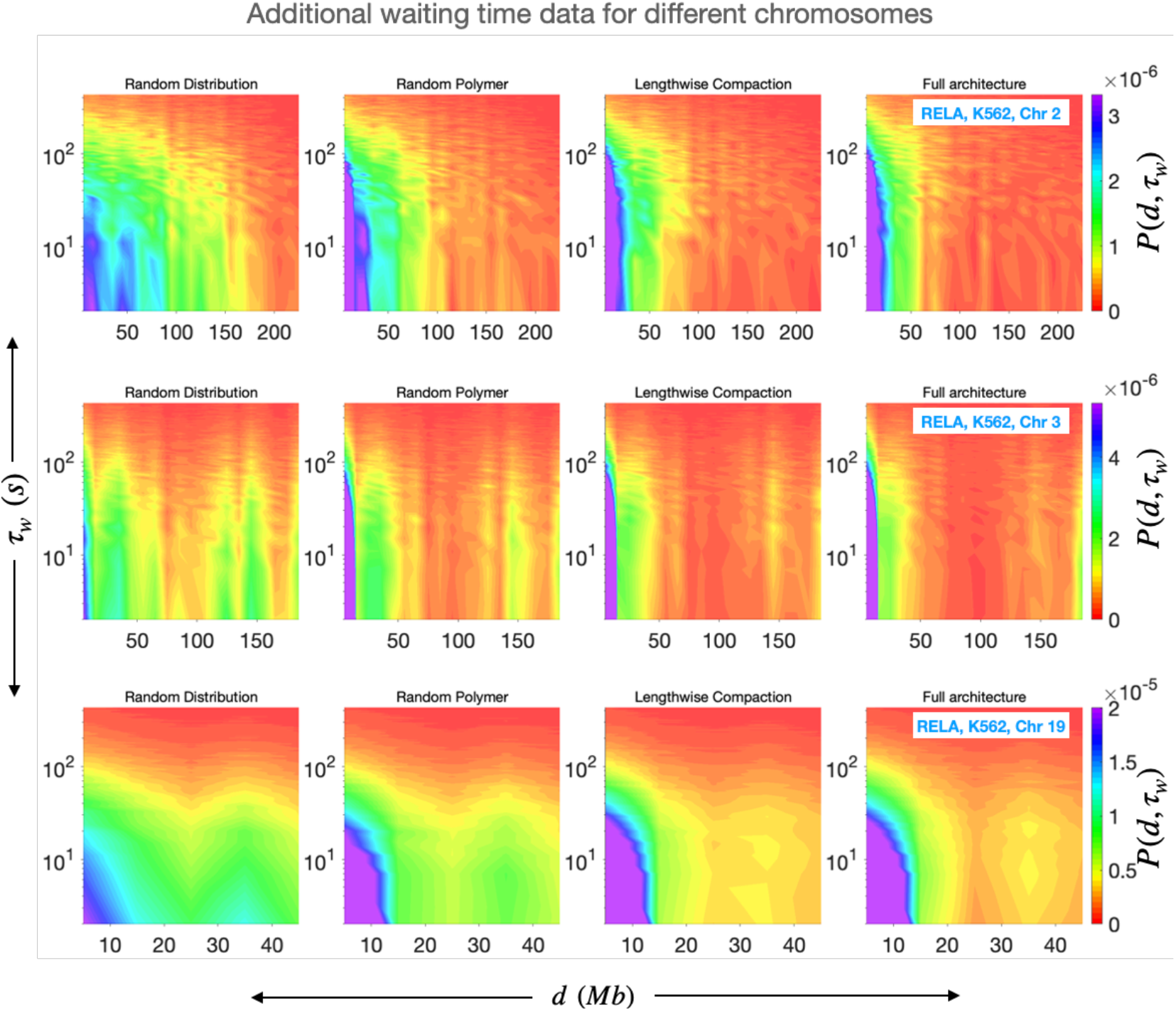
Bi-variate PDFs of waiting times and genomic distances for different chromosomes.

**FIG. S4.**
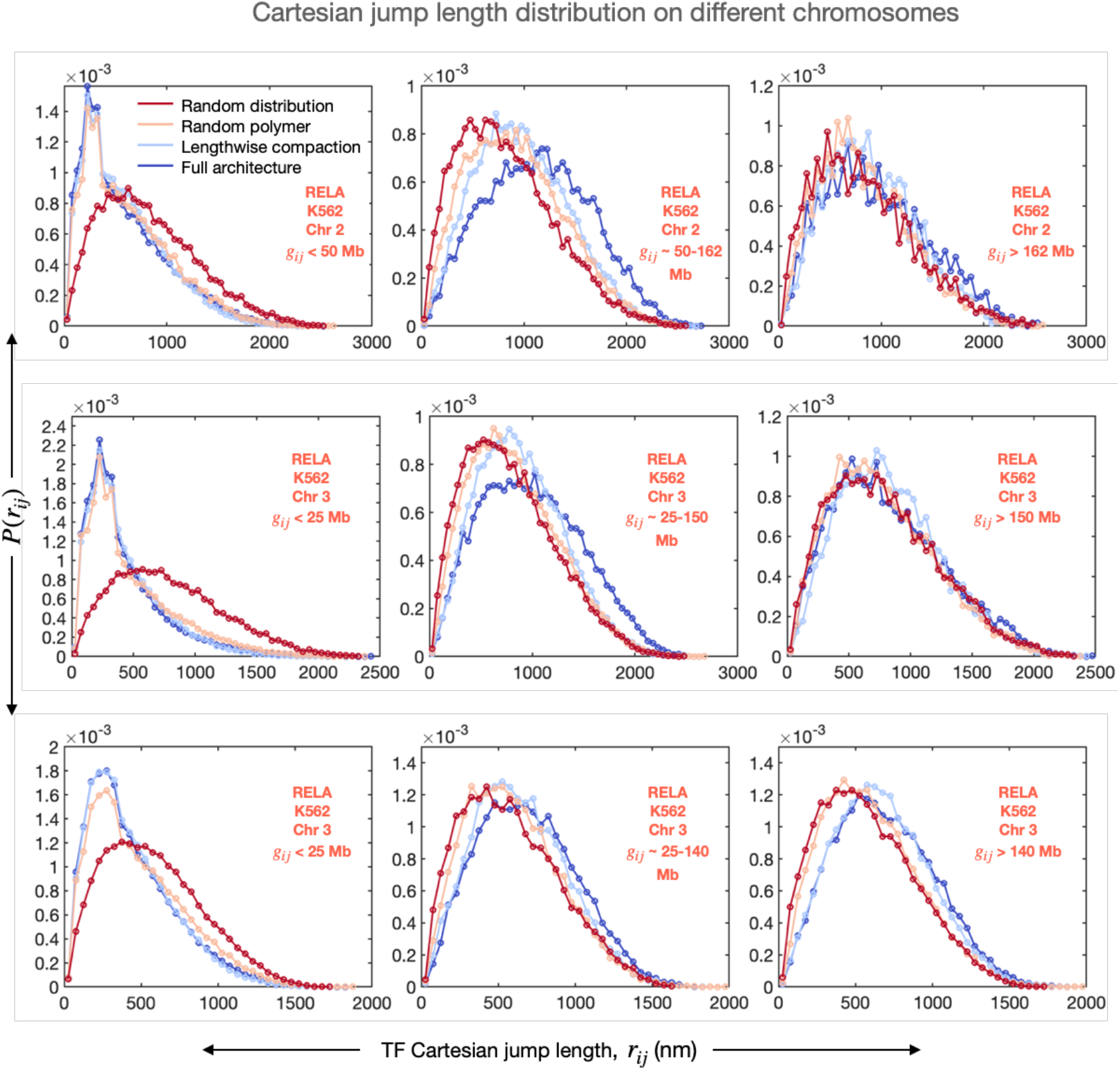
PDFs of Cartesian distances corresponding to TF commutes for three specific ranges of genomic distances for different chromosomes.

**FIG. S5.**
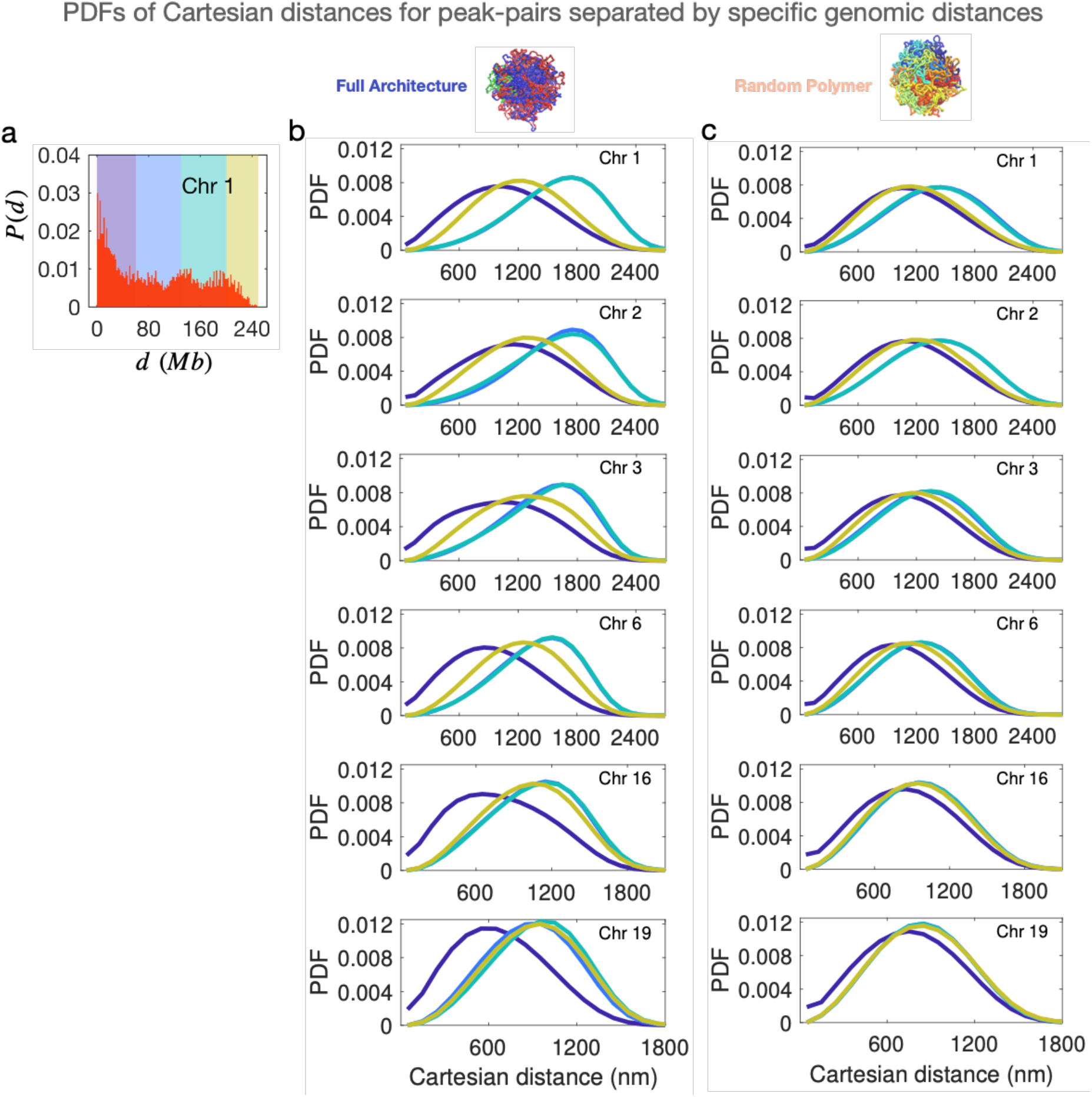
PDFs of Cartesian distances for ChIP-seq peak-pairs separated by specific genomic distances.

**FIG. S6.**
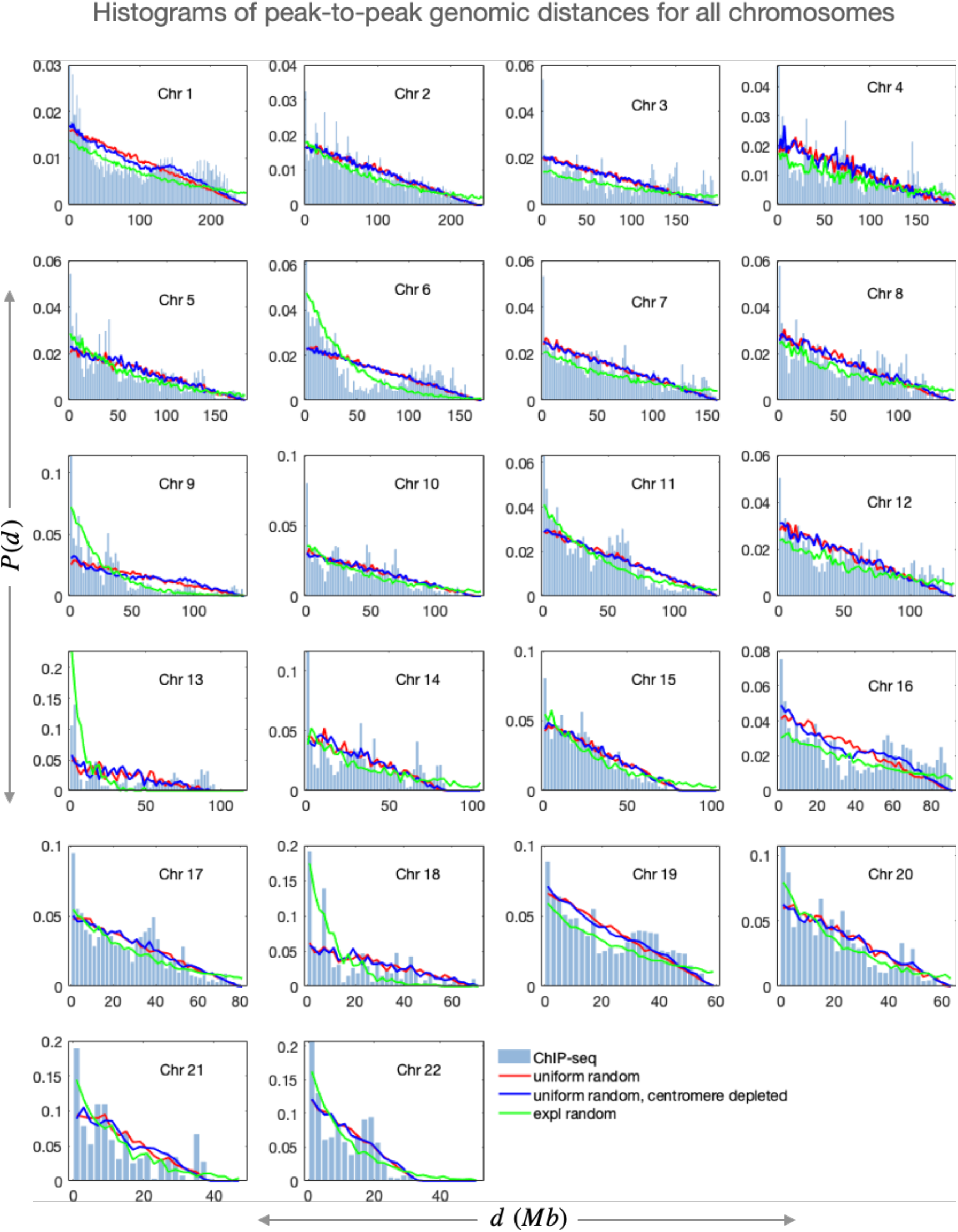
Histograms of peak-to-peak genomic distances for all chromosomes, along with predictions for various random distributions of the peak positions along the genomic sequence.

